# SuperWater: Predicting Water Molecule Positions on Protein Structures by Generative AI

**DOI:** 10.1101/2024.11.18.624208

**Authors:** Xiaohan Kuang, Zhaoqian Su, Yunchao (Lance) Liu, Xiaobo Lin, Jesse Spencer-Smith, Tyler Derr, Yinghao Wu, Jens Meiler

**Author notes:** Equal contribution (Xiaohan Kuang, Zhaoqian Su).

## Abstract

Water molecules play a significant role in maintaining protein structural stability and facilitating molecular interactions. Accurate prediction of water molecule positions around protein structures is essential for understanding their biological roles and has significant implications for protein engineering and drug discovery. Here, we introduce SuperWater, a novel generative AI framework that integrates a score-based diffusion model with equivariant graph neural networks to predict water molecule placements around proteins with high accuracy. SuperWater surpasses existing methods, delivering state-of-the-art performance in both crystal water coverage and prediction precision, achieving water localization within 0.3 ± 0.06 °A of experimentally validated positions. We demonstrate the capabilities of SuperWater through case studies involving protein hydration, protein-ligand binding, and protein-protein binding sites. This framework can be adapted for various applications, including structural biology, binding site prediction, multi-body docking, and water-mediated drug design.

## 1 Introduction

Water is essential for life, with most biological processes occurring in aqueous environments [1]. It profoundly influences the structure, stability, dynamics, and function of biomolecules within it [2, 3, 4, 5]. During protein folding, water facilitates hydrophobic interactions, drawing non-polar residues together, while also participating in hydrogen bond networks and modulating interactions between polar residues [6, 7, 8]. In protein-protein and protein-ligand interactions, water molecules not only compete with ligands for binding sites but also play a crucial role in ligand dissociation [9, 10, 11, 12]. Overlooking the role of water molecules in the process of molecular recognition has led to failures in structure-based drug discovery campaigns [13]. Thus, precise modeling of water-biomolecule interactions is crucial for effective protein design and structure-based drug development.

Despite the crucial role of water molecules, accurately determining their positions remains challenging due to the limitations of current techniques. Experimental methods often suffer from inadequate spatial and temporal resolution [3], making it difficult to fully characterize the hydration structures across entire protein surfaces [14]. To address these limitations, computational approaches, such as Molecular Dynamics and Monte Carlo simulations, have been widely employed [15, 16, 17, 18, 19]. However, these physics-based methods are heavily dependent on accurate potential energy functions and become increasingly inefficient when applied to large complexes due to their computational intensity.

With advancements in deep learning, several 3D Convolutional Neural Network (CNN) methods have been developed to predict the placement of water molecules around proteins [20, 21, 22, 23]. These methods represent the 3D structure of a protein as a 3D image, with input channels corresponding to different atom types on 3D grid points. The models are trained to differentiate between water-occupied and unoccupied voxels near the protein surface, followed by a physics-based postprocessing step to refine predictions. These 3D-CNN methods have demonstrated superior water coverage compared to traditional physics-based approaches [20, 21]. However, despite these improvements, traditional CNN-based models still encounter significant challenges. Achieving sub-angstrom accuracy in water position prediction demands very fine grid spacing, which drastically increases computational costs [24]. Additionally, these CNN-based models are sensitive to the orientation of the input structure. Although techniques like local grids or data augmentation with random rotations can alleviate this issue, they do not completely solve the problem of rotational invariance [25, 26].

In recent years, diffusion models have emerged as powerful generative AI tools [27, 28], leading to significant advancements across various areas of bioinformatics. [29, 30, 31, 32, 33, 34, 35]. Building on this foundation, we introduce SuperWater, a novel generative AI approach that leverages a score-based diffusion model combined with equivariant graph neural networks to accurately predict the placement of water molecules around proteins. Rather than directly approximating the probability distribution of water molecules around the proteins, our model learns to estimate the gradient of water distributions. This learned gradient is then used to generate water positions from a normal distribution, which are further refined using a confidence model and a clustering algorithm to optimize accuracy. SuperWater surpasses existing methods, delivering state-of-the-art results in both the coverage of crystal water molecules and the precision of predicted positions. This approach holds promise for a wide range of applications, including structural studies, binding site predictions, multi-body docking, and water-aided drug design.

## 2 Methods

### 2.1 Overview

The SuperWater pipeline comprises three main stages. The process begins with the preprocessing of the protein structure and nearby small molecules or metal atoms into heterogeneous geometric graphs, formatted for model processing. These graphs are then fed into a score-based diffusion model to sample potential positions for water molecules around the protein structure. After the initial sampling, an equivariant graph neural network-based confidence model assigns scores to each candidate water position, filtering out low-confidence predictions. Finally, a clustering algorithm consolidates neighboring water molecule positions into a single representative point for each cluster.

Fig. 1 provides a visual illustration of the entire workflow. The primary steps of SuperWater include:

**Fig. 1:**
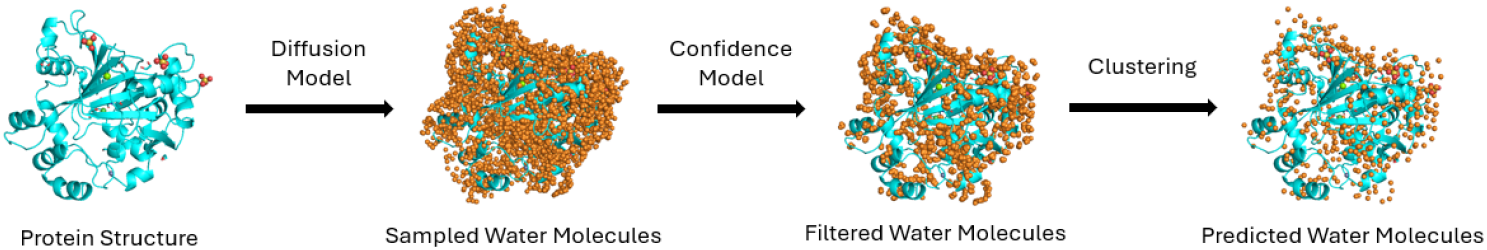
SuperWater Workflow.

**Fig. 2:**
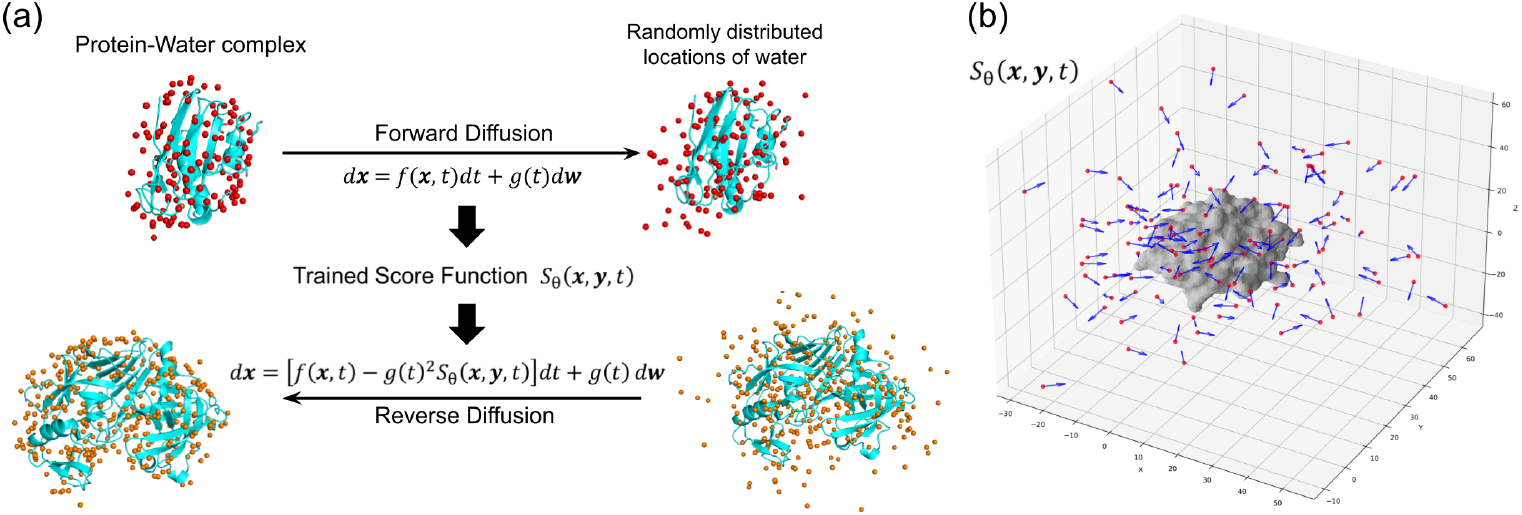
Overview of the diffusion process in the protein-water complex. (a) The forward diffusion process adds noise to generate a random distribution of water molecule positions around the protein, while the reverse diffusion leverages the trained score function *S*_*θ*_(**x, y**, *t*) to guide these positions towards energetically favorable locations. (b) Visualization of the score function *S*_*θ*_(**x, y**, *t*) as a vector field, with the protein surface shown in gray and water molecules represented by red dots. The blue arrows indicate the direction of diffusion, illustrating how the score function directs water molecules toward optimal binding sites around the protein.

- A diffusion model that operates on geometric graphs of the protein structure, allowing for the sampling of water molecule placements around the protein.
- An equivariant graph neural network-based confidence model that assesses each sampled location, discarding low-confidence predictions.
- A post-processing step that enhances prediction accuracy by clustering the predicted positions.

### 2.2 Datasets and Preprocessing

We collected high-resolution protein structures (resolution better than 1.5 °A) from the PDBBank [36], resulting in an initial dataset of 23,189 PDB files. To analyze the water binding modes, we examined all protein pairs with sequence similarity greater than 90%, as illustrated in Fig. S1. Our analysis revealed that even proteins with high sequence similarity (*>* 90%) exhibit distinct root-mean-square deviation (RMSD) values and variations in water distribution. This finding suggests that water molecules can adopt different binding modes or positions, even in highly similar protein sequences. Consistent with other studies that have clustered datasets based on binding modes rather than sequence similarity [37, 38, 39], we retained all selected PDB structures to capture the diversity of water-binding interactions and enhance model generalizability. To further refine the dataset, we selected protein structures containing between 100 and 500 residues.

Water molecules within a 4 °A radius of the protein were retained, as these fall within the second solvation shell and are involved in hydrogen bonding and hydrophobic interactions [40, 41, 42, 43, 44]. Each water molecule was represented solely by its oxygen atom[20]. For structures with multiple models, such as those with alternate residue conformations, only the first model was used. To ensure an adequate representation of water molecules, we retained only structures with a water-to-residue ratio greater than 0.6, yielding a final dataset of 17,092 PDB files. This dataset was randomly split into training, validation, and testing sets in an 8:1:1 ratio, resulting in 13,674 structures for training, 1,709 for validation, and 1,709 for testing.

### 2.3 Score-based Diffusion Model

In the diffusion model, each data point represents the 3D coordinates of a water molecule’s position on the protein structure. The generative model seeks to estimate the probability distribution *p*(**x** | **y**), where **x** indicates water positions and **y** denotes the protein structure [31]. Estimating *p*(**x**| **y**) poses two significant challenges.

The first challenge arises from the intractability of directly computing the probability distribution, as it requires normalizing the distribution across the entire space of possible positions. To address this, rather than estimating *p*(**x** | **y**) directly, the diffusion model estimates its gradient, ∇_**x**_ log *p*(**x** | **y**), known as the score function, **S**_*θ*_(**x** | **y**) [28, 45]. The score function **S**_*θ*_(**x** | **y**) is a vector field that directs water molecules toward favorable configurations from their current positions in 3D space.

The second challenge arises from the lack of sufficient training data in certain regions of the protein. To address this, the true data distribution is ‘evolved’ into a known distribution, typically a normal distribution [27, 46]. Through this process, the model diffuses data (i.e., water molecules) across the entire three-dimensional space conditioned on the protein structure, effectively filling in knowledge gaps and learning from a broader range of information across the protein structure. The entire architecture and technical details are depicted in the Supporting Information.

#### 2.3.1 Forward Diffusion and Training with SDE

The score-based diffusion model utilized in this study leverages stochastic differential equations (SDE) to generate the three-dimensional coordinates for water molecule positions around protein structures. The forward diffusion process transforms the data distribution *p*(**x**) into a Gaussian reference distribution *p*_*T*_ (**x**) through the gradual addition of noise. The dynamics of this process are governed by an SDE:

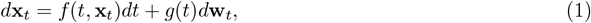

where **x**_*t*_ represents the noisy data at time *t, f* (*t*, **x**_*t*_) is the drift coefficient, *g*(*t*) is the diffusion coefficient, and **w**_*t*_ is a Wiener process. This formulation allows for a continuous transition from the real data distribution to a noise distribution, typically Gaussian.

The noise schedule is controlled by the variance *σ*_*t*_, which evolves over time according to:

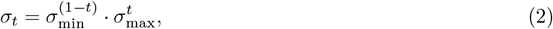

where *σ*_min_ and *σ*_max_ are the minimum and maximum noise scales, respectively. The resulting SDE can then be expressed in terms of the time-evolving variance:

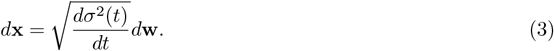

To approximate *p*(**x**_0_|**x**_*t*_), the reverse diffusion process is also modeled using an SDE:

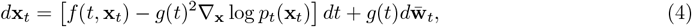

where ∇_**x**_ log *p*_*t*_(**x**_*t*_), also known as the score function, guides the denoising process. The reverse-time SDE is parameterized by a neural network, which learns to predict the score function, effectively guiding the data back towards the original, noise-free distribution.

In the specific case of predicting the positions of water molecules around proteins, the score function can be represented as:

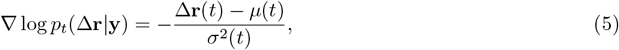

where Δ**r**(*t*) is the displacement at time *t, µ*(*t*) is the mean displacement, and *σ*^2^(*t*) represents the variance at time *t*. This formulation helps guide the denoising process by indicating the direction and magnitude in which the noisy data should be adjusted.

During training, the model optimizes a loss function to learn the score function. The primary loss is defined as:

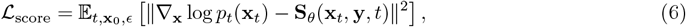

where **S**_*θ*_(**x**_*t*_, **y**, *t*) is the score function predicted by the neural network parameterized by *θ*. The current model was trained for 300 epochs using this loss function to optimize its performance by ensuring that it accurately learns to approximate the true gradient of the data distribution.

#### 2.3.2 Reverse Diffusion and Sampling

In the reverse diffusion process, the objective is to transform noisy data back into the original data distribution by reversing the effects of the forward diffusion. This process is modeled using a stochastic differential equation (SDE), similar to the forward process but in reverse time. The reverse-time SDE is given by:

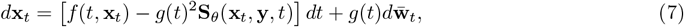

where *f* (*t*, **x**_*t*_) represents the drift coefficient, *g*(*t*) is the diffusion coefficient, and **S**_*θ*_(**x**_*t*_, *t*) is the score function predicted by the model, which guides the denoising process. The term 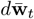 is a reverse Wiener process that models the stochastic component of the reverse diffusion.

The reverse diffusion process relies on the score function, which is learned during training. The score function **S**_*θ*_(**x**_*t*_, *t*) provides the direction and magnitude of the adjustments needed to progressively denoise the data, effectively reversing the noise added during the forward process. The model parameterized by *θ* predicts the score function and guides the sampling process to recover realistic data.

To generate samples, the model starts from the final noisy state **x**_*T*_ (which follows a Gaussian distribution) and iteratively applies the reverse SDE to gradually remove noise and restore the original structure. The step-wise application of the reverse-time SDE allows the model to produce realistic water molecule positions around the protein structure.

For each protein structure, the inference process of the diffusion model generates candidate water molecule positions, with the number of candidates being 15 times the number of protein residues. These candidates are initially distributed randomly in space based on the maximum noise level. The candidate positions are then guided by the learned score function, **S**_*θ*_(**x**_*t*_, **y**, *t*), ultimately converging to their most favorable positions within the protein structure.

### 2.4 Confidence Model

The confidence model in SuperWater is based on SE(3)-equivariant convolutional networks, adapted from the previously developed diffusion model architecture [31]. The primary purpose of this model is to score and filter the water molecule positions sampled during the inference process. Specifically, the confidence model assigns a score to each sampled water molecule by evaluating its distance to the nearest crystal water molecule [20].

The training objective is to minimize the mean squared error (MSE) between the predicted confidence scores, *p*, and the normalized distances, *N* (*d*):

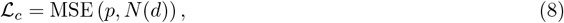

where the normalized distance *N* (*d*) is defined as:

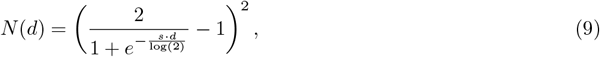

Here, *d* represents the distance between a sampled water molecule and its nearest crystal water molecule, and *s* is a scaling factor. The normalization function *N* (*d*) ensures that the distances fall within the [0, 1] range, facilitating smoother learning during training [20]. This normalization helps make the confidence scores interpretable and allows differentiation between high and low-confidence predictions.

Incorporating the normalized distances *N* (*d*) within the loss function allows the model to learn confidence scores that reflect the proximity of sampled water molecules to true crystal water positions. Higher confidence scores correspond to smaller distances, indicating greater accuracy. This is crucial for postprocessing, as it enables effective filtering of low-confidence positions, ensuring optimal placement of water molecules in the protein structure.

### 2.5 Clustering Mechanism

The final step involves refining the predicted positions of water molecules. Initially, the predicted water molecule coordinates are filtered based on their confidence scores, retaining only those with confidence scores above a defined threshold. Next, the pairwise distance matrix of the water molecule coordinates is calculated to identify neighboring water molecules that are within the van der Waals radius of oxygen, which is 1.52 °A. For each water molecule, its confidence score is compared with those of its neighbors; if it has the highest confidence among its neighbors, it is designated as the core of a cluster, and its neighbors are added to that cluster.

For each cluster, the water molecule position is refined using a confidence-weighted average, ensuring that molecules with higher confidence have a greater influence on the final position of the cluster. If a cluster contains only one water molecule, the position of that molecule is directly taken as the cluster center. After the initial clustering, a second round of filtering is performed to prevent clashes between water molecules. The pairwise distances between cluster centroids are recalculated, and clusters with distances less than 1.52 °A are further screened to retain only the centroid with the highest confidence, while removing others in close proximity. The final output includes the refined water molecule positions along with their associated confidence scores, ensuring that the predicted water molecule locations are both accurate and unique.

### 2.6 Evaluation

To assess the accuracy of predicted water molecule positions, we utilized three primary metrics: precision, coverage (recall), and root mean square deviation (RMSD).

Precision is defined as the ratio of true positive water molecule positions (TP) to the total number of predicted positions, which includes both true positives and false positives (FP). True positives (TP) are predicted water molecule positions that correctly match the experimentally determined positions within a distance cutoff of 1 °A or 0.5 °A, while false positives (FP) are predicted positions that do not have any corresponding experimentally determined water molecule within the cutoff. Precision measures how many of the predicted positions are actually correct:

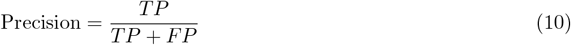

Coverage, also referred to as recall, represents the proportion of true water molecule positions that were correctly predicted by the model. Specifically, coverage is calculated after matching at most one crystal water to each predicted water within a distance cutoff of 1 °A or 0.5 °A, It is defined as the fraction of matched crystal water positions among all crystal water positions:

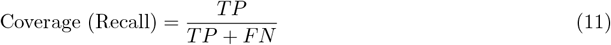

RMSD quantifies the average distance between predicted water molecules and their closest experimentally determined water positions. It is calculated as:

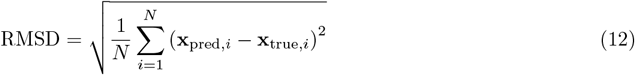

where *N* is the number of predicted water molecules, **x**_pred,*i*_ represents the position of the *i*-th predicted water molecule, and **x**_true,*i*_ is the corresponding experimentally determined position. A lower RMSD value indicates that the predicted water positions are closer to the true positions, thereby reflecting higher accuracy.

Together, these metrics provide a comprehensive evaluation of the accuracy of predicted water molecule positions. Precision assesses how many of the predicted water molecules are correct, coverage measures how effectively all true water molecules are identified, and RMSD evaluates the positional accuracy of the predictions.

## 3 Results and Discussion

This section presents a comprehensive evaluation of SuperWater, comparing its performance with the state-of-the-art HydraProt method [20] for predicting water molecule positions around protein structures. Both methods were tested on the same dataset to ensure a fair comparison. In the inference phase of SuperWater, an initial set of water molecule positions was sampled randomly throughout the system, with the number of water molecules set to 15 times the number of protein residues to maximize coverage while balancing computational cost and accuracy. These initial positions were refined through a reverse diffusion process to adjust their translational degrees of freedom. Next, a confidence model was used to filter out low-confidence positions, retaining those most likely to correspond to true water molecule locations. Finally, a clustering algorithm was applied to further optimize and finalize the predicted water molecule positions. The dataset used for evaluation consisted of protein structures curated from the Protein Data Bank (PDB). Further details regarding dataset curation, preprocessing, and experimental conditions can be found in Methods.

### 3.1 Comparison between SuperWater and HydraProt

Fig. 3 illustrates the precision versus coverage curves for SuperWater and HydraProt across different thresholds. Predictive models typically exhibit a trade-off between precision and coverage, which aligns with our observations in these figures. The curves are generated by varying the prediction thresholds for both SuperWater and HydraProt.

**Fig. 3:**
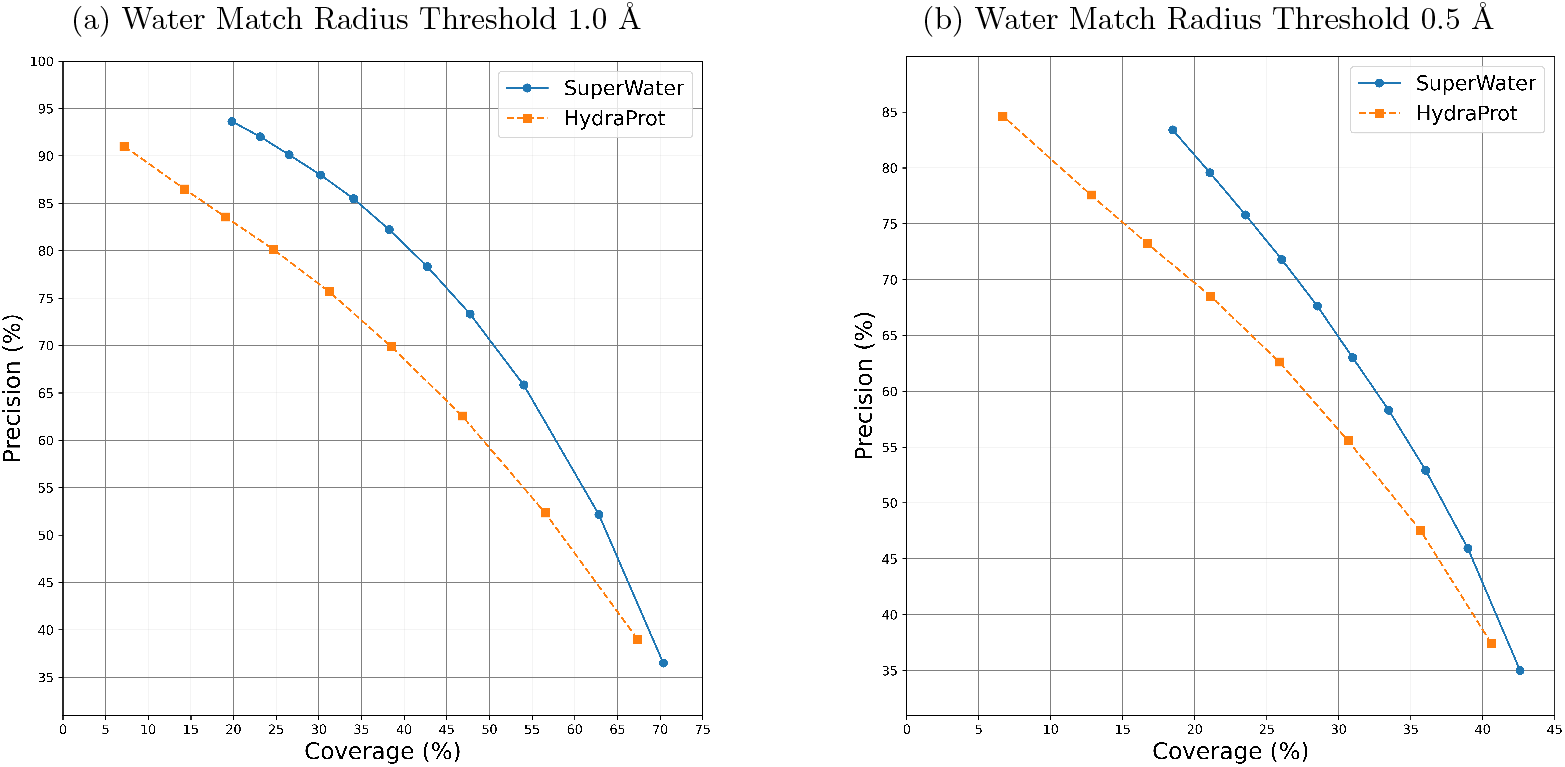
Comparison of Precision and Coverage for SuperWater and HydraProt at Different Thresholds. The graphs compare the performance of SuperWater (blue line) and HydraProt (orange line), showing how precision varies with coverage at different match radius thresholds: (a) 1.0 °A and (b) 0.5 Å.

Overall, SuperWater consistently demonstrates higher precision across a broader range of coverage levels compared to HydraProt, highlighting its superior predictive capabilities. In Fig. 3a, at a match radius threshold of 1.0 Å, SuperWater achieves 90% precision at approximately 27% coverage, whereas HydraProt reaches 90% precision at only about 8% coverage. This shows that SuperWater’s coverage is more than three times that of HydraProt at this level of precision. At a broader coverage of 54%, SuperWater maintains a precision of around 66%, significantly outperforming HydraProt, which drops to 55% precision. These results emphasize SuperWater’s ability to deliver high precision while achieving greater coverage, enhancing the reliability and scope of its predictions.

At a match radius threshold of 0.5 Å (Fig. 3b), SuperWater achieves approximately 70% precision at 27% coverage, while HydraProt reaches only about 61% precision at the same coverage level. As coverage increases to 40%, SuperWater maintains an precision of around 46%, whereas HydraProt’s precision drops to 37%. These findings underscore SuperWater’s superior performance across varying cutoff criteria.

Beyond precision and coverage in predicting water-binding sites, it is also important to assess the spatial accuracy of these predictions. To achieve this, we calculated the RMSD between the predicted water positions and their corresponding nearest crystal water positions, as shown in Fig. 4. The graph reveals that SuperWater consistently yields lower RMSD values compared to HydraProt, demonstrating higher spatial accuracy across different precision thresholds. Further analysis of mean absolute deviation (MAD) between experimentally determined water positions and the correctly predicted positions (true positives) is provided in the supplementary material. At a probability threshold of *cap* = 0.5, SuperWater achieves a MAD of 0.3 ± 0.06 Å, demonstrating the robustness of its predictions. However, in real-world applications, the actual positions of predicted water molecules are generally unknown. Therefore, we employ RMSD, calculated between all predicted positions and their nearest experimental counterparts, as a comprehensive metric to assess the model’s overall spatial prediction performance.

**Fig. 4:**
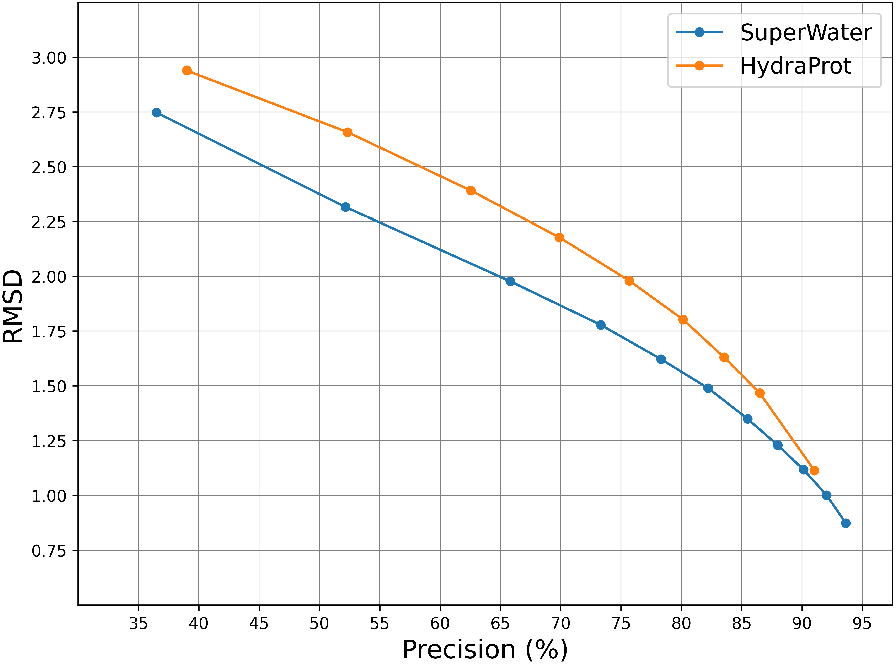
Comparison of Root Mean Square Deviation (RMSD) between SuperWater (blue) and HydraProt (orange) Across Different precision Levels. RMSD measures the average spatial deviation between all predicted water molecule positions and experimentally determined positions, serving as a metric for the overall spatial precision of predictions. Lower RMSD values indicate higher spatial accuracy in the predicted positions.

### 3.2 Case Studies

To assess the performance and biological relevance of SuperWater, we present three case studies that showcase its capabilities in accurately predicting conserved water molecule positions across various scenarios: protein surfaces, protein-ligand binding sites, and protein-protein interaction interfaces, as depicted in Fig. 5, Fig. 6, and Fig. 7, respectively. The figures were generated using PyMOL[18], with experimental water molecules depicted in red, SuperWater-predicted water molecules in cyan, and HydraProt-predicted water molecules in yellow.

**Fig. 5:**
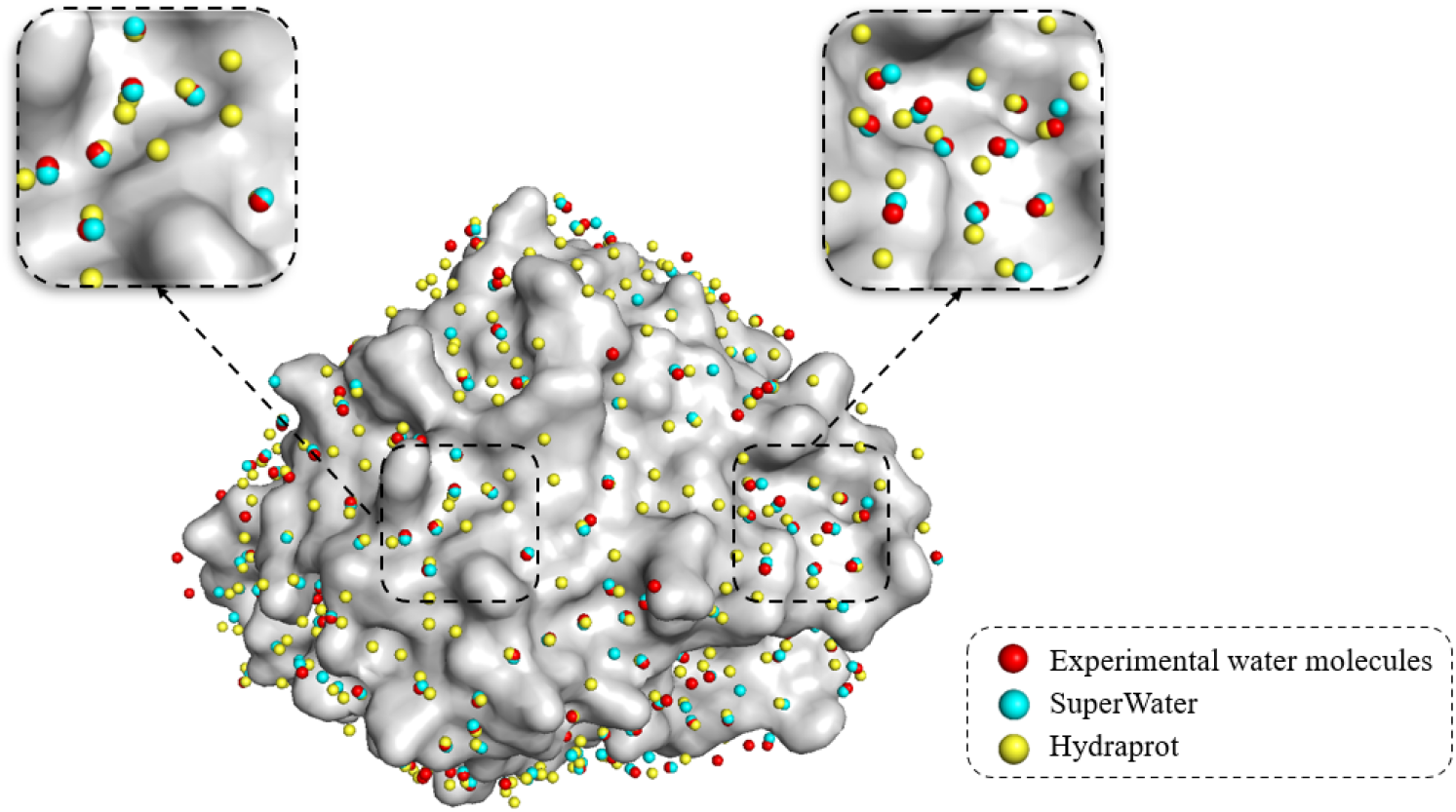
Visualization of experimental water molecules (red), SuperWater predicted water molecules (cyan), and Hydraprot predicted water molecules (yellow) around Carbonic Anhydrase II (PDB ID: 6OUH).

**Fig. 6:**
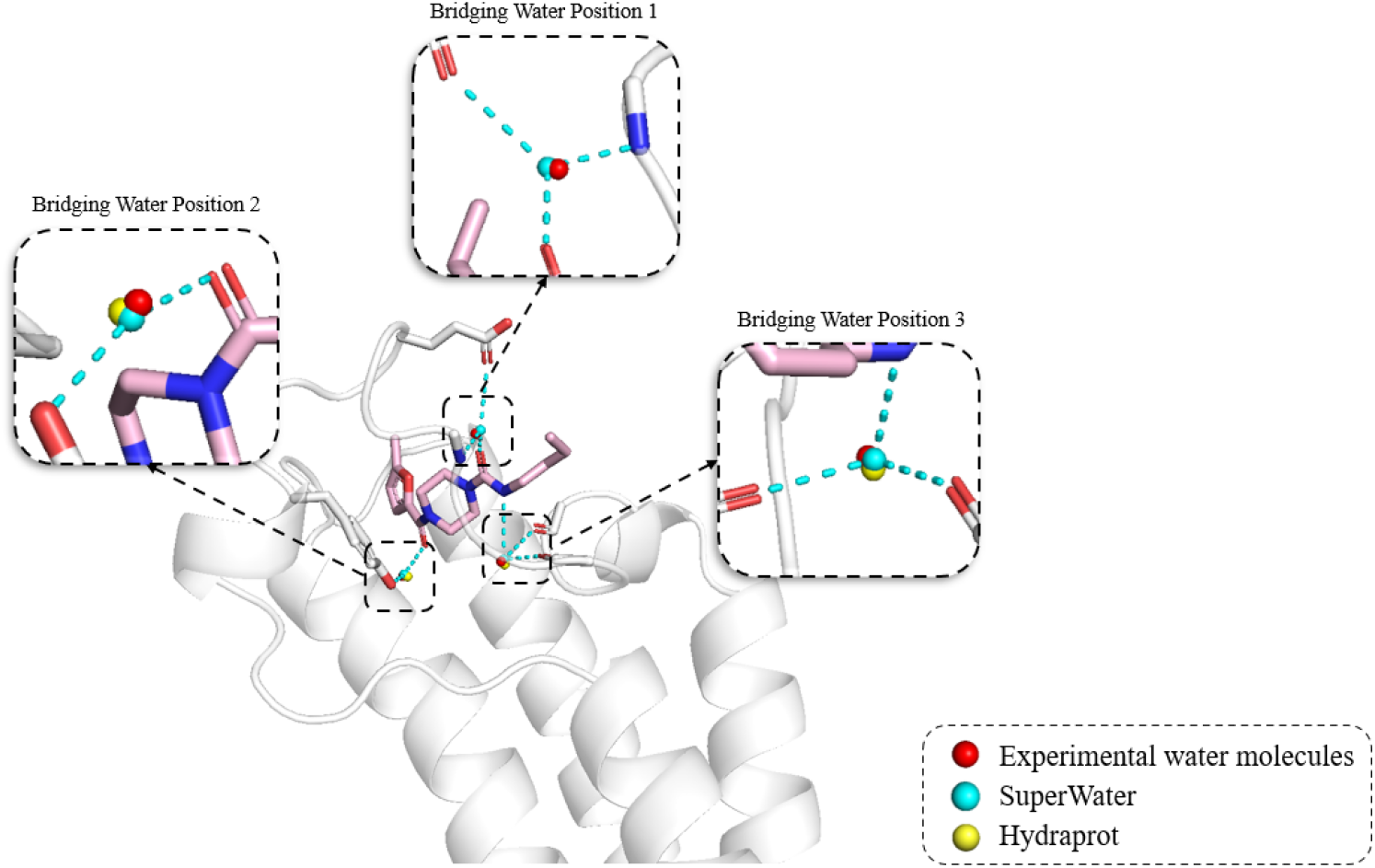
Visualization of experimental water molecules (red), SuperWater-predicted water molecules (cyan), and HydraProt-predicted water molecules (yellow) around the PHIP protein (PDB ID: 7FVP). Three panels zoom into each of the three bridging water positions, which are labeled as Bridging Water Position 1, Bridging Water Position 2, and Bridging Water Position 3. Notably, SuperWater successfully captures all three bridging water positions, whereas HydraProt fails to identify the bridging water at Position 1.

**Fig. 7:**
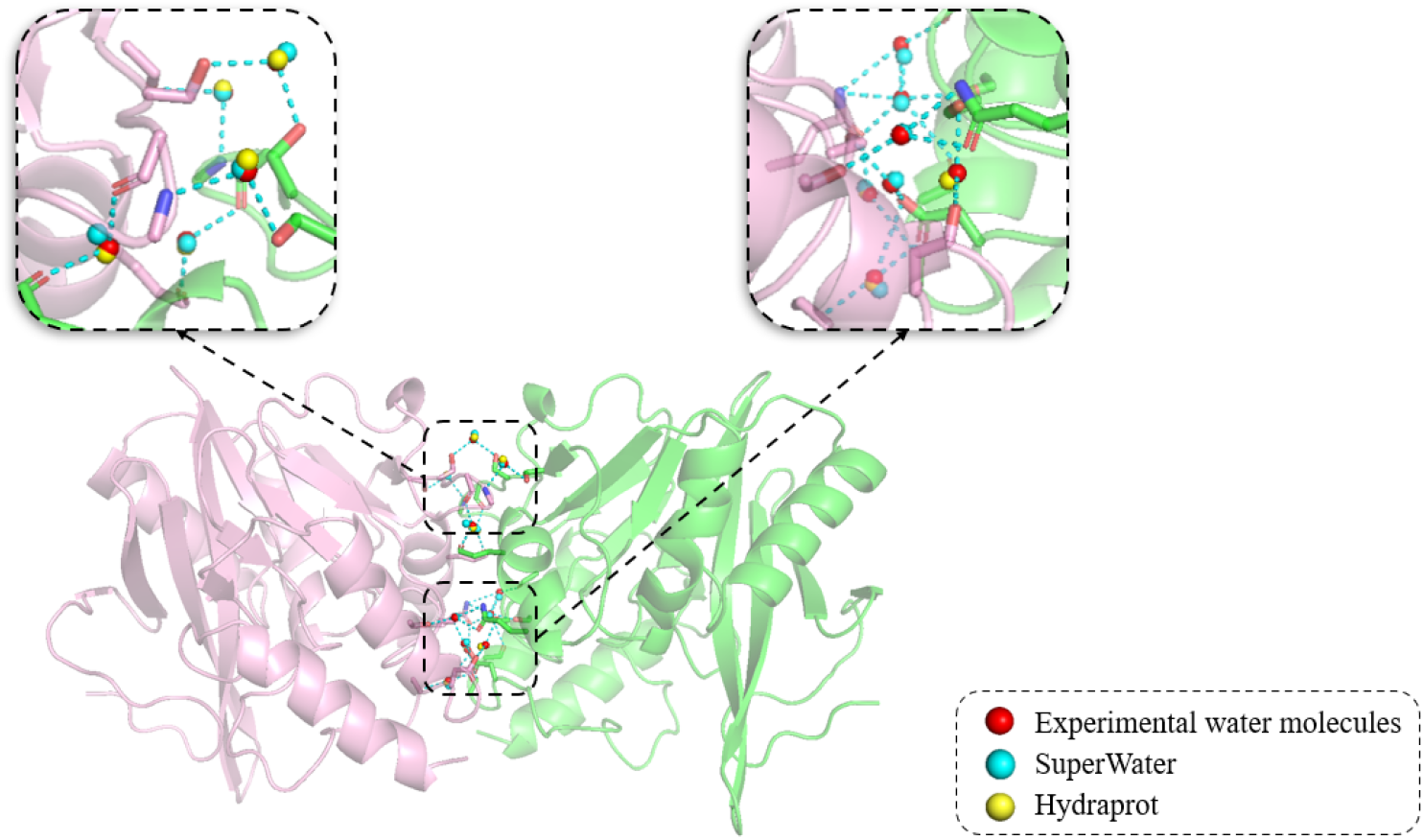
Visualization of experimental water molecules (red), SuperWater predicted water molecules (cyan), and Hydraprot predicted water molecules (yellow) of protein-protein binding sites (PDB ID: 4EY2).

#### 3.2.1 Protein Surfaces

The water molecules on proteins are necessary for their folding, stability, and functions [47]. Fig. 5 illustrates the conserved water molecule predictions on the surface of Carbonic Anhydrase II (PDB ID: 6OUH). The SuperWater-predicted water molecules (cyan) align closely with experimentally observed positions, with minimal false positives, indicating high precision. In contrast, HydraProt-predicted water molecules (yellow) are dispersed across a broader spatial region, resulting in reduced prediction precision due to numerous misplaced water molecules.

Table 1 summarizes the computational evaluation metrics for SuperWater and HydraProt on the 6OUH structure. SuperWater consistently outperforms HydraProt across recall, precision, F1 score, and mean absolute deviation (MAD) at thresholds of both 0.5 Å and 1.0 Å. The superior performance of SuperWater, particularly in terms of precision and F1 score, underscores its capability to accurately predict water molecule positions on protein surfaces.

**Table 1:** Comparison of Computational Evaluation Metrics for SuperWater and Hydraprot for 6OUH

| Metrics<br>Models | Recall |  | Precision |  | F1 Score |  | MAD (Å) |  |
| --- | --- | --- | --- | --- | --- | --- | --- | --- |
|  | 0.5 | 1.0 | 0.5 | 1.0 | 0.5 | 1.0 | 0.5 | 1.0 |
| SuperWater cap 0.05 | **0.761** | **0.851** | **0.620** | **0.693** | **0.683** | **0.764** | **0.241** | **0.288** |
| HydraProt cap 0.05 | 0.475 | 0.707 | 0.262 | 0.390 | 0.338 | 0.503 | 0.262 | 0.418 |

#### 3.2.2 Protein-Ligand Binding Sites

In this case study, we evaluate SuperWater’s ability to predict bridging water molecules within the binding pocket of the PHIP protein (PDB ID: 7FVP). Bridging water refers to water molecules that simultaneously form hydrogen bonds with both the protein and the ligand. These molecules play a crucial role in stabilizing the structure of the complex and facilitating the ligand recognition process [9].

Fig. 6 provides a detailed comparison of the experimental, SuperWater-predicted, and HydraProt-predicted water positions. In the binding pocket, three key bridging water molecules are highlighted, labeled as Bridging Water Position 1, Position 2, and Position 3. SuperWater accurately predicted all three bridging water positions, whereas HydraProt missed the crucial bridging water at Position 1.

Table 2 summarizes the computational evaluation metrics for SuperWater and HydraProt for all experimental waters in the 7FVP structure. The results indicate that SuperWater consistently outperforms HydraProt in recall, precision, F1 score, and mean absolute deviation (MAD) at thresholds of 0.5 Å and 1.0 Å. Notably, SuperWater’s superior precision and F1 score highlight its capability to accurately predict water molecule positions that are essential for maintaining complex stability and supporting water-mediated drug design.

**Table 2:** Comparison of Computational Evaluation Metrics for SuperWater and Hydraprot for 7FVP

| Metrics | Recall |  | Precision |  | F1 Score |  | MAD (Å) |  |
| --- | --- | --- | --- | --- | --- | --- | --- | --- |
| Models | 0.5 | 1.0 | 0.5 | 1.0 | 0.5 | 1.0 | 0.5 | 1.0 |
| SuperWater cap 0.05 | **0.705** | **0.855** | **0.581** | **0.705** | **0.637** | **0.773** | **0.231** | **0.313** |
| HydraProt cap 0.05 | 0.487 | 0.674 | 0.325 | 0.450 | 0.390 | 0.539 | 0.287 | 0.403 |

#### 3.2.3 Protein-Protein Interaction Interfaces

Water molecules are also essential for mediating protein-protein interactions. Accurately predicting interfacial water molecules is essential for understanding protein interactions, which is vital for designing inhibitors or stabilizers for protein complexes. Fig. 7 highlights the bridging water molecules at the protein-protein interface of the NDM1-meropenem complex (PDB ID: 4EY2). The experimental structure reveals 12 bridging water molecules at this interface. SuperWater predicts 11 of these water molecules that form hydrogen bonds between the two protein chains with high precision, whereas HydraProt predicts only 8, with their positions falling within a 1 Å radius of the actual experimental water molecules. Further details regarding these bridging water molecules can be found in Supplementary Figs. S6 and S7.

Table 3 summarizes the computational evaluation metrics for SuperWater and HydraProt for all experimental waters in the 4EY2 structure. The results indicate that SuperWater consistently outperforms HydraProt in terms of recall, precision, and F1 score at thresholds of 0.5 Å and 1.0 Å. However, HydraProt shows a slightly better mean absolute deviation (MAD) than SuperWater in this case. Accurately predicting interfacial water molecules is essential for understanding protein interactions, which plays a critical role in designing effective inhibitors or stabilizers for protein complexes.

**Table 3:** Comparison of Computational Evaluation Metrics for SuperWater and Hydraprot for 4EY2

| Metrics<br>Models | Recall |  | Precision |  | F1 Score |  | MAD (Å) |  |
| --- | --- | --- | --- | --- | --- | --- | --- | --- |
|  | 0.5 | 1.0 | 0.5 | 1.0 | 0.5 | 1.0 | 0.5 | 1.0 |
| SuperWater cap 0.05 | **0.496** | **0.706** | **0.361** | **0.516** | **0.418** | **0.596** | 0.285 | 0.410 |
| HydraProt cap 0.05 | 0.485 | 0.704 | 0.250 | 0.362 | 0.330 | 0.479 | **0.265** | **0.398** |

## 4 Conclusion

In this study, we presented SuperWater, a novel generative AI model designed for accurately predicting the positions of water molecules around protein structures. Our approach combines a score-based diffusion model with an equivariant graph neural network to generate candidate water placements, evaluate their confidence, and refine the predictions. The performance of SuperWater was benchmarked against the state-of-the-art HydraProt model.

SuperWater demonstrated superior performance across multiple metrics, including precision, coverage, and overall spatial accuracy. Specifically, SuperWater outperformed HydraProt by achieving higher precision at broader coverage levels. For example, at a precision level of approximately 90%, SuperWater achieved nearly three times the coverage compared to HydraProt, highlighting its ability to identify a larger set of correct water positions without compromising precision. Furthermore, SuperWater consistently produced lower RMSD values, showcasing its spatial precision and robustness in predicting water locations. In all three case studies, SuperWater outperformed HydraProt in terms of recall, and precision, effectively capturing key water-mediated interactions in protein surfaces, protein-ligand binding, and protein-protein interfaces. These results underscore SuperWater’s robustness and versatility, establishing it as a valuable tool for accurate water placement predictions on protein structures. Reliable water position prediction is crucial for understanding water-mediated biological processes, protein stability, and lead optimization in drug discovery. Although SuperWater is currently trained to predict water molecule positions, its framework can be readily adapted for diverse applications, including structural biology, metal-binding site prediction, multi-body docking, and water-mediated drug design.

## Data and Code Availability

The processed datasets and cache files are available on Zenodo (https://doi.org/10.5281/zenodo.14166655). The source code for SuperWater is available on GitHub (https://github.com/kuangxh9/SuperWater).

## Supporting Information

Supplementary information is available.

## Acknowledgements

Z.S. thanks the support of the Vanderbilt Data Science Postdoctoral Fellowship. X.L. and X.K. are grateful for the research funding and support provided by the Vanderbilt Data Science Institute. Y.L acknowledges the Nvidia hardware grant for accelerating the project development. X.L. also thanks the John R. Hall Professorship Endowment in Chemical Engineering for its support. J.L. expresses gratitude for the project opportunity provided by the Vanderbilt Data Science Institute. We sincerely thank Umang Chaudhry for facilitating access to these resources. The authors thank Tommi Jaakkola and Gabriele Corso at Massachusetts Institute of Technology for the help and guidance. We also acknowledge the computational resources (DGX A100) provided by the Vanderbilt Data Science Institute. J.M. is supported by a Humboldt Professorship of the Alexander von Humboldt Foundation. J.M. acknowledges funding by the Deutsche Forschungsgemeinschaft (DFG) through SFB1423 (421152132), SFB 1664 (514901783), TRR (514664767), and SPP 2363 (460865652). J.M. is supported by the Federal Ministry of Education and Research (BMBF) through the Center for Scalable Data Analytics and Artificial Intelligence (ScaDS.AI), through the German Network for Bioinformatics Infrastructure (de.NBI), and through the German Academic Exchange Service (DAAD) via the School of Embedded Composite AI (SECAI 15766814). Work in the Meiler laboratory is further supported through the National Institute of Health (NIH) through R01 HL122010, R01 DA046138, R01 AG068623, U01 AI150739, R01 CA227833, R01 LM013434, S10 OD016216, S10 OD020154, S10 OD032234.

## Author contributions

Z.S. conceptualized and designed the study and prepared the dataset. Z.S., X.K., Y.L., X.L., J.S., T.D., Y.W., and J.M. contributed to software development, model training, and analysis. Z.S., X.K., Y.L., J.S., T.D., Y.W., and J.M. contributed to the writing and review of the manuscript. J.S., J.M., and T.D. provided funding support.

## Notes

### Competing Interest Statement

The authors have declared no competing interest.

